# A multiplexed homology-directed DNA repair assay reveals the impact of ~1,700 BRCA1 variants on protein function

**DOI:** 10.1101/295279

**Authors:** Lea M. Starita, Muhtadi M. Islam, Tapahsama Banerjee, Aleksandra I. Adamovich, Justin Gullingsrud, Stanley Fields, Jay Shendure, Jeffrey D. Parvin

## Abstract

Loss-of-function mutations in *BRCA1* confer a predisposition to breast and ovarian cancer. Genetic testing for mutations in the *BRCA1* gene frequently reveals a missense variant for which the impact on the molecular function of the BRCA1 protein is unknown. Functional BRCA1 is required for homology directed repair (HDR) of double-strand DNA breaks, a key activity for maintaining genome integrity and tumor suppression. Here we describe a multiplex HDR reporter assay to simultaneously measure the effect of hundreds of variants of BRCA1 on its role in DNA repair. Using this assay, we measured the effects of ~1,700 amino acid substitutions in the first 302 residues of BRCA1. Benchmarking these results against variants with known effects, we demonstrate accurate discrimination of loss-of-function versus benign variants. We anticipate that this assay can be used to functionally characterize BRCA1 missense variants at scale, even before the variants are observed in results from genetic testing.

## Introduction

Genetic testing for hereditary breast and ovarian cancer genes often reveals a missense variant in BRCA1 whose impact on the molecular function of the encoded protein, and therefore its contribution to cancer risk, is unknown. These variants are reported as “variants of uncertain significance” (VUS). VUS cause distress for both physicians and patients and can lead to unnecessary surgeries^1,2^. As genetic testing becomes more common in the clinic, reports of missense VUS are rapidly accumulating in public databases^3^. Additional VUS arise from the increasingly widespread sequencing of tumor genomes and exomes to guide precision therapy. Therapeutic agents, such as PARP inhibitors, that specifically target BRCA-deficient tumors, are an effective therapy only for the subset of tumors with specific types of *BRCA1* mutation [for example refs. 4,5]. Therefore, knowledge of germline and somatic *BRCA1* mutation status is important to guide both cancer prevention and treatment strategies.

The most commonly reported class of VUS in *BRCA1* are single nucleotide variants (SNVs) that are predicted to result in missense amino acid substitutions. Currently, 1,794 missense VUS in BRCA1 are in the clinical genetics database Clinvar^3^. An additional 218 missense variants have conflicting interpretation reports, suggesting that clinical testing labs apply discordant classification criteria. Across BRCA1’s 1863 amino acids, there are 12,458 SNVs that are predicted to result in a missense substitution that might or might not affect protein function; these become VUS when identified as a germline or somatic variant in a patient. Most strategies for variant interpretation fail when the variants are sufficiently rare^6–9^. Computational variant-effect prediction algorithms scale without limit, but these are not accurate enough for routine clinical use^10^. Functional assays, on the other hand, are considered by the American College of Medical Genetics (ACMG) guidelines as strong evidence for or against the pathogenicity of missense variants^10^. However, performing a *post hoc* functional assay for each BRCA1 SNV as it is discovered is infeasible at their current rate of accumulation.

BRCA1 is required for maintenance of genome integrity via the homology-directed DNA repair (HDR) pathway. The effect of variants in BRCA1 on its HDR function can be determined in tissue culture using a GFP-based reporter assay for intact DNA repair function^11^. For BRCA1 variants tested thus far, this assay has stratified the protein function of known benign and pathogenic variants with high sensitivity and specificity^12–15^. Here, we describe a multiplex version of this HDR assay that we developed toward the goal of testing all possible protein variants in the N-terminus of BRCA1 (residues 2-302), which includes the RING domain (residues 7-98). Proper folding of the RING domain is required for the stability and function of the full-length protein^13,16,17^. In addition, missense mutations that cause increased cancer risk frequently map to either the RING or BRCT domains.

Using the HDR reporter cell line with integrated *BRCA1* variant libraries, we test approximately 600 BRCA1 missense variants per experiment. Damaging mutations are identified by their relative depletion from the subset of cells that are GFP-positive. We show that, as expected, the first 98 amino acids of BRCA1 which encode the RING domain are more sensitive to substitutions than the subsequent 204 residues. The results of the multiplexed assay correlate well with the results from singleton HDR reporter and other functional assays. Furthermore, known pathogenic variants can be separated from known benign variants by their recurrent depletion from the functional population in across replicate experiments. Our results for 66 VUS or variants with conflicting reports of pathogenicity from ClinVar show that seven are nonfunctional for HDR. We suggest improvements to our current protocol that could increase its throughput and accuracy. Finally, we anticipate that this assay can be used to functionally characterize BRCA1 missense variants at scale in order to provide the additional information necessary to more definitively interpret VUS in the clinic.

## Results

### A multiplexed assay to measure the effect of protein variants on homology-directed DNA repair

The multiplexed HDR reporter assay is based on an assay developed by the Jasin laboratory^11^. In this reporter assay, a site-specific double-stranded DNA break induced by the I-SceI endonuclease results in conversion of a GFP-negative cell to GFP-positive if the HDR pathway is intact (Fig. 1a). Loss of BRCA1 activity (*e.g.* by depletion with siRNA) results in cells that cannot repair the GFP through HDR from the donor template. We previously used this assay^13–15,18^ to test the capacity of 158 individual BRCA1 missense variants to rescue the loss of endogenous BRCA1. BRCA1 HDR function with these individual assays demonstrates 100% specificity and 91.6% sensitivity for predicting the cancer risk associated with variants with established interpretations (n = 43, Supplementary Fig. 1). The one known pathogenic variant misidentified as benign, R71G, affects splicing, rather than protein function^19^, and therefore could not have been classified correctly with an assay that expresses variants within a cDNA copy of *BRCA1*. Given this functional assay’s established specificity and sensitivity for predicting clinical pathogenicity, we sought to convert it to a multiplexed format in order to generate high-confidence predictions for hundreds of BRCA1 variants in a single experiment.

**Figure 1.**
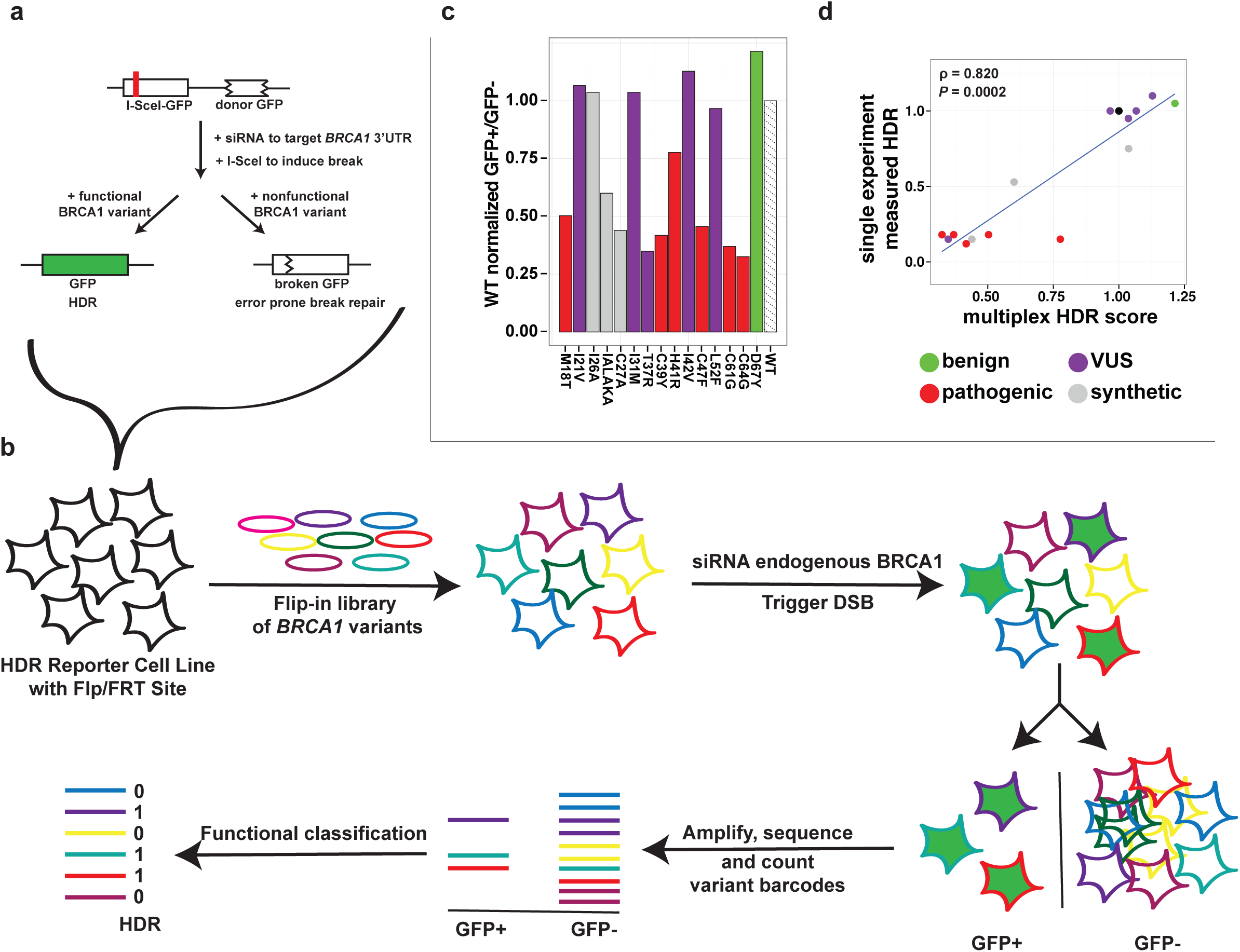
Overview of the multiplexed-HDR reporter assay. **a,** Schematic of the integrated HDR reporter. **b,** Schematic of workflow for multiplexed HDR-reporter assay. **c,** Results from the pilot 16-plex HDR assay testing WT and 15 variants in a multiplexed format. WT-normalized, GFP-positive:GFP-negative ratios are plotted on the y-axis with variant identifications on the x-axis. **d,** The correlation between scores from the 16-plex experiment and scores from individual HDR-reporter assays. Spearman rho and p-value are reported. Bar and points are colored according to ClinVar variant interpretation.

As a pilot version of a multiplexed HDR reporter assay (Fig. 1b), we prepared a pool of plasmids containing the wild-type (WT) *BRCA1* and 15 *BRCA1* variants that had been tested individually. We integrated these plasmids *en masse* into the HeLa-derived HDR reporter cell line, which we had engineered to contain a single recombinase-based landing pad (HeLa-DR-FRT, Methods). Though it is low efficiency, we used a recombinase-based system to ensure that only a single variant would be integrated into a common site in each cell, ensuring consistent expression levels and avoiding complications associated with lentiviruses, *e.g.* variable integration sites and template switching^20^.

After selecting for cells containing an integrated *BRCA1* variant, we performed the three steps of the HDR reporter assay. First, endogenous BRCA1 was depleted with an siRNA targeting the 3’ untranslated region of the endogenous *BRCA1* mRNA. Second, I-SceI was expressed to generate a double-strand break in the broken GFP reporter. Third, after allowing sufficient time for double-stranded DNA break repair, cells were sorted into GFP-positive and GFP-negative populations. The *BRCA1* variants in each sorted population were PCR amplified and sequenced, and DNA reads for each variant were counted. We calculated a score for each variant by taking the ratio of its frequency in the GFP-positive cells to its frequency in the GFP-negative cells, and normalized to the equivalently calculated ratio for the WT transgene. Scores from this pilot assay were largely consistent with published results for these variants (Fig. 1c, d); the sole exception was the BRCA1 H41R variant, which had an intermediate level of activity in the multiplex assay. Known pathogenic mutants were defective in this pilot experiment and the single known benign missense variant had similar repair activity to the wild-type BRCA1. We therefore proceeded to scale up the assay to analyze hundreds of variants per experiment, focusing on BRCA1 residues 2-302.

We created three individual pools of barcoded site-saturation mutagenesis libraries^14,21^. Pool1 contains variants in residues 2-96, pool2 in residues 97-192, and pool3 in residues 193-302 (Supplementary Table 1). We integrated the plasmids in each pool into the HeLa-DR-FRT cell line. We then performed four replicates of the multiplexed HDR reporter assay on each pool, using either an siRNA against endogenous *BRCA1* mRNA or a control siRNA (Supplementary Table 2). The barcodes were amplified and sequenced from the genome of the cells in the GFP-positive and GFP-negative populations. The score for each variant was calculated as described above for the small-scale assay (**Supplementary Table 4**). Variant scores between replicates were well correlated for pool1, but less so for pools 2 and 3, for which most variants scored close to ln(0), indicating no depletion (Supplementary Fig. 2)

### Functional classification of variants

A characteristic of all sequencing count data, including from multiplexed assays for variant effect, is that the variance of scores increases at low read counts due to error from Poisson-distributed shot noise and stochastic dropout of variants^22^. This effect is exacerbated in cases like the HDR reporter assay that, despite being the gold standard assay for *BRCA1* missense variants, have a bottleneck. Specifically, only ~10% of the cells at most become GFP-positive; this bottleneck limits the dynamic range of functional scores (Supplementary Table 2). Because the score depends on read count, we cannot directly compare the scores of any two variants. Therefore, we chose to construct a binary classifier (“depleted “vs. “not depleted”) by modeling the relationship between the read count and score (Methods). With this model, we tested the significance of each variant’s depletion in the *BRCA1* siRNA experiments compared to variants in the control siRNA experiments at the same read count in the same experiment (Fig. 2a, Supplementary Fig. 3, Methods, and **Supplementary Table 4**). To remove as many variants as possible that could be substantially affected by stochastic dropout, we applied a stringent read count filter to the GFP negative population and removed variants below this threshold from further analysis (Supplementary Fig. 3, Methods).

**Figure 2.**
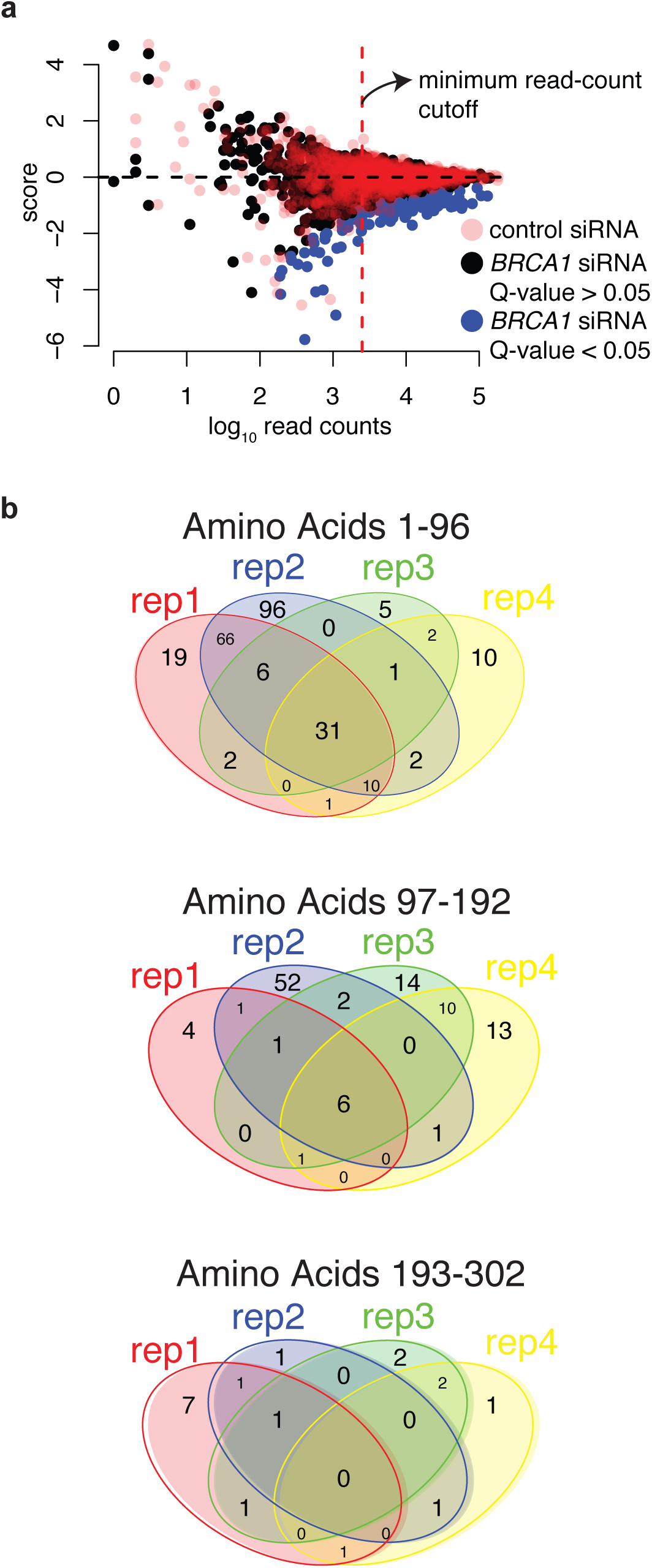
Identifying BRCA1 variants depleted from the GFP-positive population. **a,** The log of the WT-normalized, GFP-positive:GFP-negative ratios are on the y-axis and log_10_ read counts are on the x-axis for a single replicate of the HDR-reporter assay for codons 2-96. Variants from the control (pink) or BRCA1 (black) siRNA conditions are indicated, variants significantly depleted from the GFP-positive population in the *BRCA1* siRNA condition, q < 0.05, colored blue. The dashed line represents the read-count threshold. **b,** Venn diagrams of the number of variants found depleted in multiple or single replicates.

For each experimental replicate, the number of variants above the read-count threshold in the GFP negative population varied, as did the number of variants classified as depleted (*i.e.* nonfunctional variants, Table 1). The Venn diagrams shown in Fig. 2b indicate the number of variants found to be depleted among the 4 replicate experiments for each pool of variants. In each set of replicate experiments, some variants were scored as depleted only once. Most of these singletons were depleted due to stochastic dropout, but some were nonsense variants that passed the read-count filter only in one replicate (**Supplementary Table 4**). The variants found to be depleted in 3 or 4 replicates were enriched in nonsense codons and residues with known pathogenic mutations (Fisher’s exact test, 2X enrichment, p = 2.8 x 10^-6^ and 20X enrichment, p = 2.0 x 10^-29^, respectively). Pool1 had the highest percentage of nonfunctional variants, which was expected because the structured RING domain is found almost entirely within the amino acids mutated in pool1. Pool2 ranked second, with substitutions at the only two remaining positions of the RING domain (97 and 98) repeatedly depleted from the functional GFP-positive population. In contrast, pool3 had relatively few depleted variants, which were mostly not shared between replicate experiments.

**Table 1.**
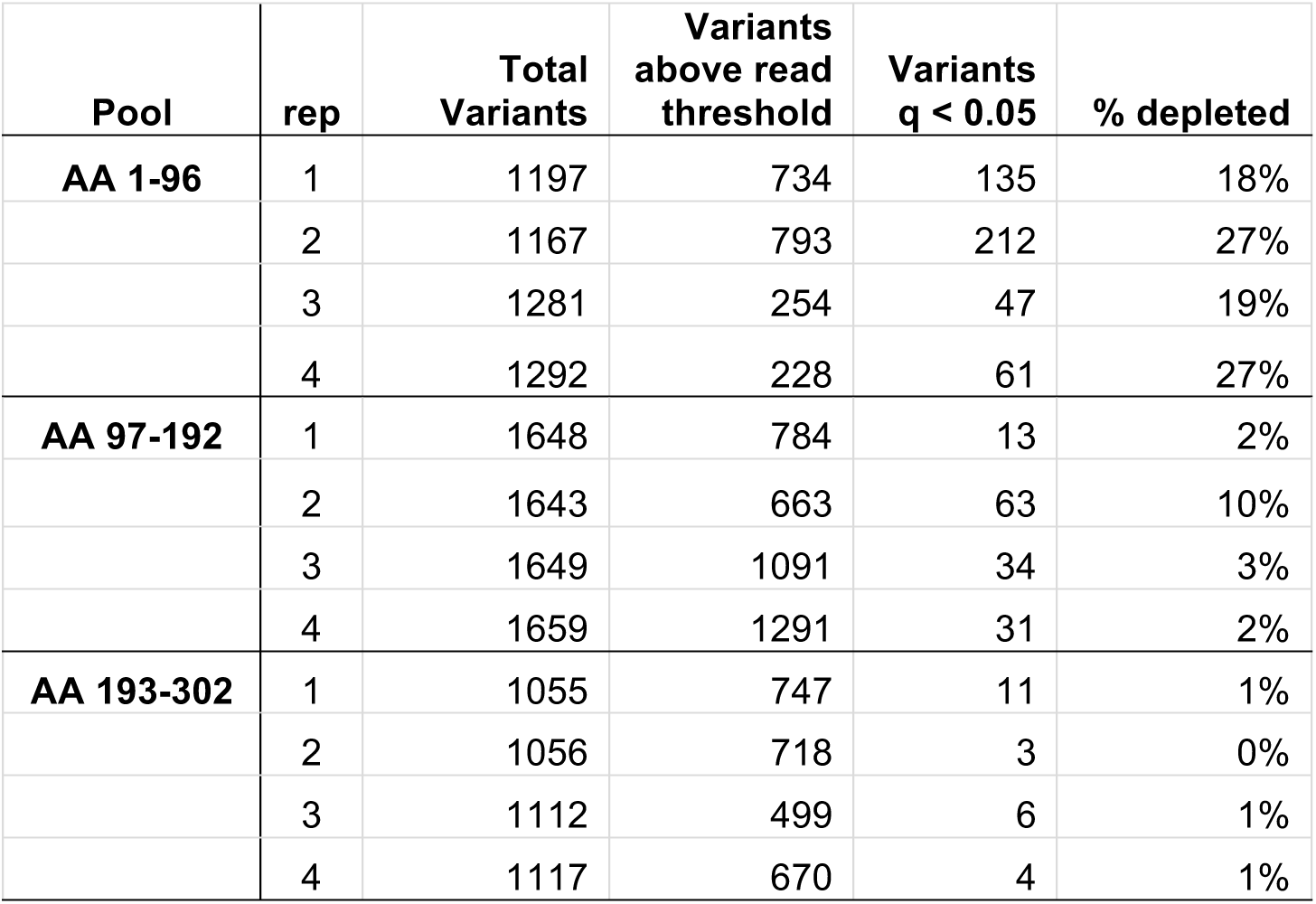

| Pool | rep | Total Variants | Variants above read threshold | Variants $q < 0.05$ | % depleted |
| --- | --- | --- | --- | --- | --- |
| AA 1-96 | 1 | 1197 | 734 | 135 | 18% |
|  | 2 | 1167 | 793 | 212 | 27% |
|  | 3 | 1281 | 254 | 47 | 19% |
|  | 4 | 1292 | 228 | 61 | 27% |
| AA 97-192 | 1 | 1648 | 784 | 13 | 2% |
|  | 2 | 1643 | 663 | 63 | 10% |
|  | 3 | 1649 | 1091 | 34 | 3% |
|  | 4 | 1659 | 1291 | 31 | 2% |
| AA 193-302 | 1 | 1055 | 747 | 11 | 1% |
|  | 2 | 1056 | 718 | 3 | 0% |
|  | 3 | 1112 | 499 | 6 | 1% |
|  | 4 | 1117 | 670 | 4 | 1% |

### Variant effects measured in this multiplexed HDR assay are strongly concordant with singleton HDR assays and ClinVar interpretations

In total, we measured the functional effect of 1,696 BRCA1 missense or nonsense variants in three or more replicates. The great majority of BRCA1 missense variants were functional for DNA repair, with only 61 variants (3.6%) depleted from the population in 3 or 4 replicates and therefore likely to be nonfunctional for DNA repair (Fig. 3a). Among the variants successfully tested, results from singleton HDR reporter assays were available for 15 variants^12,14,15^. All 11 that had been scored as functional in HDR were depleted in 0 replicates in the multiplexed assay, and all 4 that were nonfunctional in singleton assays were all depleted in 3 or 4 replicates of the multiplexed assay (Fig. 3b).

**Figure 3.**
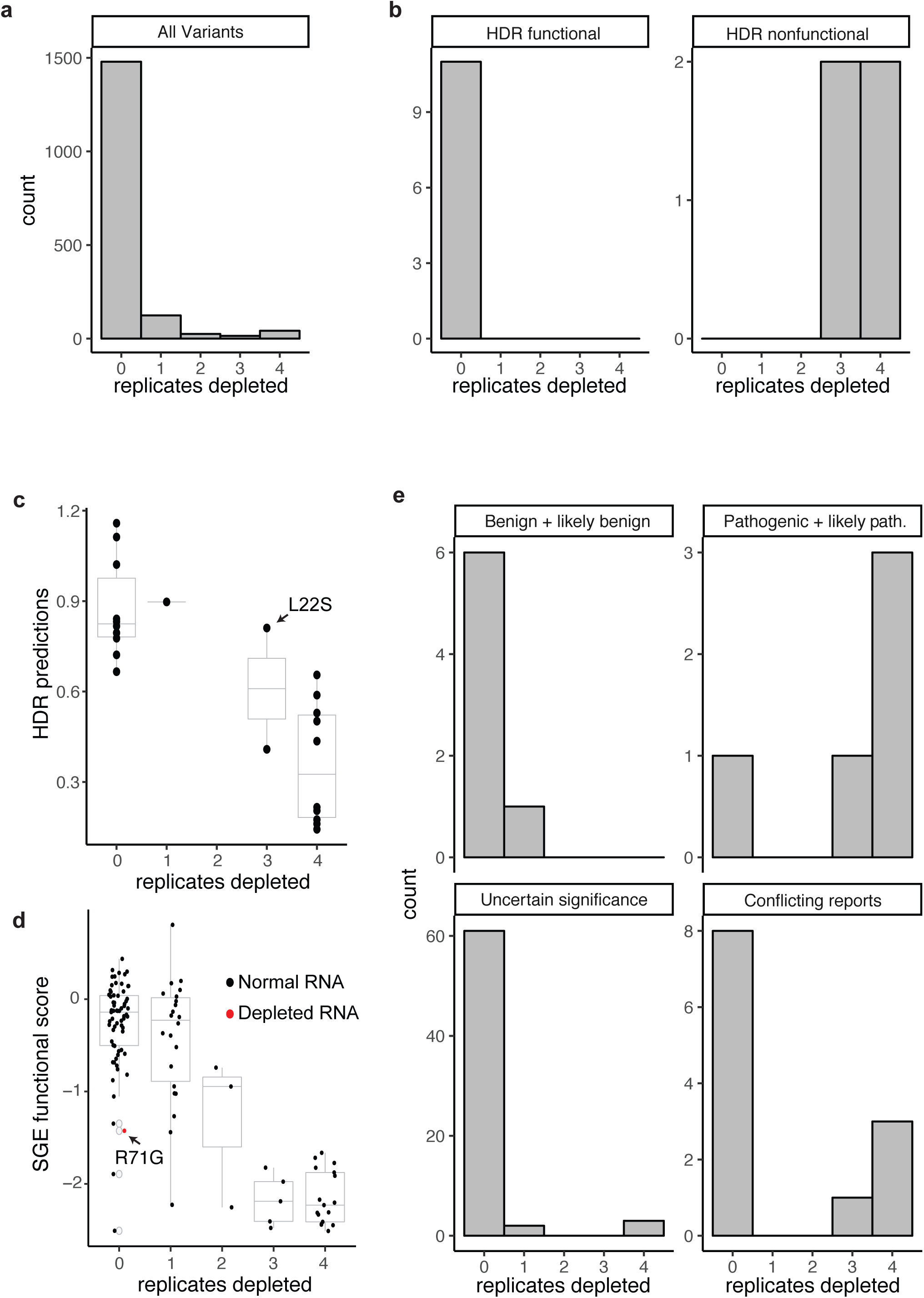
Comparison of depletion scores to scores from other functional assays and ClinVar classifications. **a,** Histogram of depletion score for all variant above the read count threshold in at least three replicates. **b,** Histograms of variant depletion scores that were functional or nonfunctional as measured by individual HDR assays. **c,** Box and strip plots comparing variant depletion scores (x-axis) to HDR predictions (y-axis) from ref.16; BRCA1 L22S is indicated. **d,** Box and strip plots comparing variant depletion scores (x-axis) to SGE functional scores (y-axis; Findlay et al, unpublished), points marking variants with >80% RNA depletion in the SGE assay are colored red; BRCA1 R71G is indicated. **e,** Histograms of variant depletion scores as for each ClinVar classification.

The number of times that variants were depleted from the functional population in the multiplexed HDR reporter assay was strongly correlated with the results of other multiplexed functional assays. Specifically, we previously used *in vitro* ubiquitin ligase and BARD1-binding yeast two-hybrid scores to predict HDR function for variants within the first 102 amino acids of BRCA1^14^. The 25 variants that overlap in the two assays are highly concordant (Fig. 3c). Variants that were never depleted in the multiplexed HDR reporter assay correspond to those in the previous study with higher HDR predictions, whereas variants that were repeatedly depleted correspond to those in the previous study with lower HDR predictions. However, the phage and yeast-based functional assays misidentified the known pathogenic mutant L22S as functional, whereas it was correctly found to be defective in both single^14^ and multiplexed HDR experiments in human cells.

The number of replicates in which variants were depleted from the multiplexed HDR reporter assay was also strongly consistent with functional scores from a multiplex assay for BRCA1 function that relies on survival of a haploid cell line after saturation genome editing (SGE) of exons in *BRCA1* (Fig. 3d; Findlay et al, enclosed). To an extent, the exceptions are predictable. The SGE-based functional assay identifies variants that reduce splicing because the variants are edited into their native context in the genome, whereas the they are false negatives in the HDR reporter assay in which the BRCA1 ORF is not spliced (Fig. 3d, red points represent variants with reduced RNA levels).

Like the multiplexed HDR reporter assay, the SGE assay is highly accurate for identifying variants that are damaging for protein function. However, there are variants that are found to be damaging to protein function by SGE and not the multiplexed HDR reporter assay (V14G, C44G, E85G, A92G, and A92T). In previous work, we have individually compared the effects of amino acid substitutions on BRCA1 function in HDR versus repair by single-strand annealing (SSA)^15^. Though HDR and SSA are both related mechanisms for repairing DNA double-strand breaks dependent on BRCA1, there were different tolerances for seven of the 35 variants tested when comparing the two assays in that study. While all seven of these were functional in HDR, three were non-functional in SSA and four had significantly depleted partial function in SSA^15^. When comparing the two multiplexed assays for HDR and SGE, finding five differences from among 1,696 measurements of BRCA1 function in HDR is a low number and may reflect appropriate biology of the two assays. This is especially true given that the biochemical mechanism affecting cell growth in the SGE assay is unknown. Of the five differences when comparing the high-throughput HDR and SGE assays, C44G may be a false negative of the multiplexed HDR reporter assay given that all amino acid substitutions in the cysteine and histidine residues that bind zinc ions in the BRCA1 RING domain that have been tested to date in singleton functional assays were damaging^12–14^, although this specific C44G variant has not before been tested for HDR function. Alternatively, C44G and the other discordant variants may be slightly destabilized and rescued by the expression level in the HDR reporter assay, or these residues may be necessary for another function of BRCA1 critical for cell survival in the SGE assay.

Clinical interpretations for some of the BRCA1 variants that we functionally score here are reported in the ClinVar database (accessed June, 2017)^20^. Of the ~1,700 variants scored here, seven were classified as benign or likely benign in ClinVar. All seven were either never depleted, or depleted in only one replicate, in the multiplexed HDR assay. In contrast, we tested five BRCA1 variants that are established pathogenic mutations in ClinVar. Of these, four were depleted in 3 or 4 replicates in the multiplexed HDR assay. The fifth pathogenic variant, R71G, had WT-like DNA repair activity in our data, a result consistent with previous HDR reporter assays^15^. R71G causes a defect in RNA splicing^18^, which as discussed above we do not expect to be detectable in our assay (Fig. 3e).

In summary, we observe strong concordance between variants depleted in 0-1 replicates in our assay and benign status in ClinVar or WT-like function in other assays; as well as between variants depleted in 3-4 replicates in our assay and pathogenic status in ClinVar or loss-of-function in other assays. We therefore concluded that the number of replicates in which a variant is found depleted is a reasonable proxy for its functionality, and term these as “depletion scores” (range 0-4, Supplementary Table 5).

### Depletion scores identify damaging BRCA1 variants

Depletion scores for all variants passing the read-count threshold in at least three replicates are shown in the form of a sequence–function map in Fig. 4a. 100% of the damaging amino acid substitutions (dark red) were observed in the first 98 amino acids which comprises the RING domain, whereas residues 99-302 strongly tended to tolerate amino acid substitutions. Nonsense mutants, indicated with the asterisk on the bottom row were for the most part non-functional. Three of these nonsense mutants were scored as functional (codons 289, 290, and 291), and we consider these to be false negatives. The distribution of variants indicates that the degenerate positions in the mutagenic oligonucleotides used to make the site-saturation mutagenesis libraries contained a much higher fraction of guanines than the 1:1:1:1 ratio of the four nucleotides specified for the synthesis (see Methods). Thus, codons containing guanine in the first or second position (particularly glycine, arginine, alanine and valine) are the most highly represented. Since the highest representation of substitutions was to glycine, we mapped the number of times each substitution to glycine was found depleted in replicate experiments to the solution structure of the BRCA1 and BARD1 RING domain dimer^23^ (Fig. 4b). Amino acids positions that were the most intolerant to substitution were either buried in the interior of the 4-helix bundle, which acts as the BARD1 interface, or in the loops that coordinate zinc ions.

**Figure 4.**
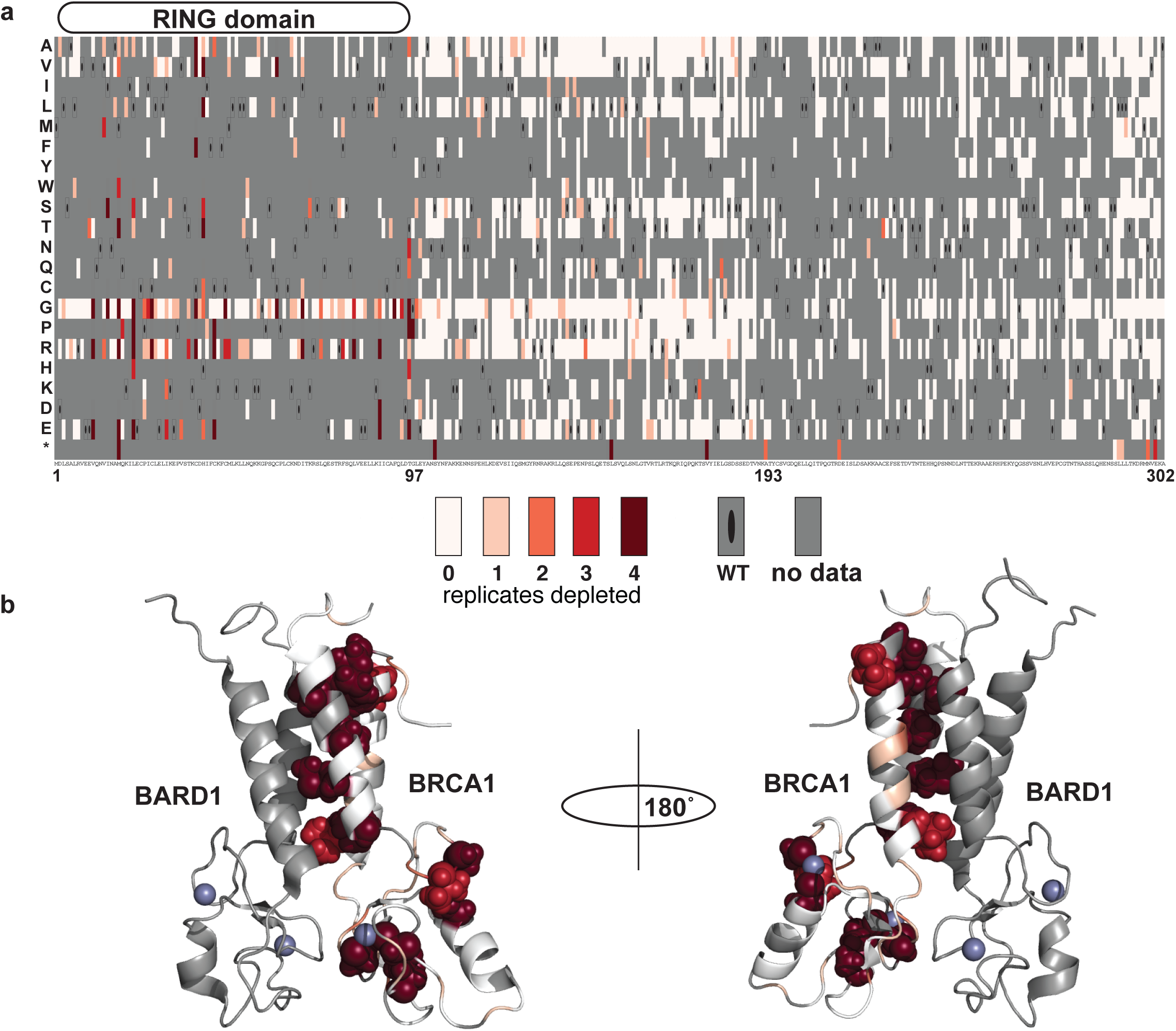
The effect of amino acid substitutions the DNA repair function of BRCA1 2-302. **a,** A sequence-function map of the effect of amino acid substitutions in BRCA1 2-302. The depletion score for each variant is the count of experimental replicates in which it was found depleted from the functional population. Each position in BRCA1(2-302) is arranged along the x-axis, the position of the RING domain is diagrammed above. The amino acid substitutions, grouped by side-chain properties, are on the y-axis, nonsense codons are *. The depletion scores range from never depleted, or likely functional (white), to likely nonfunctional in dark red. Black ovals demarcate the wild-type residue and gray, missing data. **b,** The depletion score for amino acid substitutions to glycine are mapped to the solution structure of the BRCA1 1-102, BARD1 26-125 dimer (pdb 1JM7). Color scale as in (a) and spheres are shown for side chains at amino acid postions with depletion score of 3 or 4.

We examined the depletion scores of 77 BRCA1 variants that are ambiguously classified in ClinVar (VUS or conflicting reports of pathogenicity) to predict their functional impact. Of 66 variants tested in the multiplexed HDR reporter assay that were classified as VUS in ClinVar, three (V11G, C47G and I68R) had a depletion score of 3-4 and are thus likely nonfunctional for HDR. Of 12 variants that have conflicting interpretations of pathogenicity in ClinVar, four (M18T, T37R, C39F and H41L) were nonfunctional in the DNA repair assay (Fig. 3e). These seven variants, all ambiguous in ClinVar and nonfunctional in our assay, are either at positions that coordinate zinc ions (C39F, C47G, H41L), at positions within the zinc-finger loops (T37R and I68R) or to have side chains in the interior of the 4-helix bundle (V11G and M18T). Most known pathogenic missense mutations map to these same structural features. Of note, the four variants currently listed in ClinVar as having conflicting interpretations of pathogenicity and nonfunctional here were interpreted in the past as likely pathogenic or pathogenic^3^.

The remaining 70 variants that are ambiguously classified in ClinVar (VUS or conflicting reports of pathogenicity) have a low depletion score in our assay. Of these, 60 lie outside of the RING domain where we did not identify any damaging amino acid substitutions. The 10 variants found within the structured RING include V11A, a conservative amino acid substitution inside the 4-helix bundle, and I90T, whose sidechain points out of the helix. T77M and T97A are substitutions to the amino acids that abut the helices and may not affect BARD1 binding; however, other changes at T97 are not tolerated (Fig. 4a). I42L and M48V are in the RING loops but are conservative changes and therefore may be tolerated; E29G, E33A, G57R and P58A are also in the loops, but their side chains point away from the interior of the structure which may be why they are tolerated. In studies of ubiquitin ligase activity, substitutions at E29 are damaging to ubiquitin ligase function^14^ but not to HDR. This finding adds to the growing evidence that ligase activity is not required for BRCA1’s HDR function^24,25^.

## Discussion

We developed a multiplexed reporter assay to measure the effect of hundreds of amino acid substitutions in BRCA1 on HDR, the molecular function most closely associated with its role in tumor suppression. We reproducibly measured 1,696 of the possible 6,020 amino acid or nonsense substitutions (301 × 20). Although these 1,696 constitute only 28% of all possible substitutions, our assay is the most high-throughput to date that specifically analyzes the DNA repair function of BRCA1. Of the 12 variants known to be benign or pathogenic that are assayed here, we classified them with 80% sensitivity and 100% specificity^3^. The assay was only 80% sensitive because it misclassified R71G, a mutation that is pathogenic consequent to its impact on splicing, a feature not tested in the DNA repair assay.

In this multiplex HDR reporter assay we used a cell line (HeLa) that was not derived from breast or ovarian tissue. The requirement for a high transfection efficiency restricted us to only a few choices of human cell lines. The function of BRCA1 in HDR is considered ubiquitous for all human cell types. Although the use of HeLa cells for HDR reporter assays produced results that accurately predict hereditary breast and ovarian cancer risk, this assay does not address the breast and ovarian specificity of loss of BRCA1 activity.

The non-uniform coverage of amino acid substitutions in our variant libraries was a technical challenge. Two features of the protocol employed here to make variant libraries^21^ contributed to this non-uniformity. First, inverse PCR reactions were performed individually for each amino acid position using individually synthesized oligonucleotides. Therefore, failed reactions and uneven mixing of the PCR products caused loss of all or most substitutions at some positions (Fig. 4a). Second, a strong guanine bias at each degenerate nucleotide position during oligonucleotide synthesis led to a bias in the encoded amino acids (Fig. 4a). We anticipate that if the distribution of variants in the library were more uniform, we would be able to query more variants per experiment, with a theoretical maximum of ~4,000 variants per experiment under ideal conditions, at our current number of sorted cells. Alternative approaches using array-derived oligonucleotides may create more uniformly distributed variant libraries^26,27^. It may also be useful to limit the number of amino acid changes to those accessible by single nucleotide changes, as these are the missense variants relevant to human disease. However, this limitation would also reduce the information content of the multiplexed experiments.

In addition to improving library uniformity, restricting the number of barcoded variants in each multiplexed HDR reporter experiment should result in more variants that pass the read-count filter. For example, a higher percentage of variants passed the read count threshold in pool2 (50%, on average) than pool1 (29%) and pool3 (31%). Pool3 had twice as many barcoded variants as pool2 (50,000 vs. 25,000) and covered more sequence space (106 vs. 96 amino acids positions), which resulted in nearly a 20% reduction in the number of reproducibly queried variants in pool3 compared to pool2. On the other hand, the variant libraries for pool1 and pool2 contained similar numbers of barcoded variants. However, more variants were damaging to BRCA1 HDR function in pool1, and thus only ~7% of the cells in this pool converted to being GFP-positive following the double-strand break, further limiting the dynamic range of pool 1 experiments.

In summary, we describe the development of the first multiplexed assay for measuring the effects of amino acid substitutions on a protein’s function in double-strand DNA break repair in human cells. We analyzed nearly 1,700 amino acid substitutions of BRCA1 residues 2-302 and found that this approach yielded results comparable to low-throughput HDR analysis of single variants and other functional assays. More importantly, our results are concordant with known cancer-predisposing mutants of BRCA1, including perfect positive predictive value for identifying known mutations damaging to protein function. This assay can be repurposed to measure the effect of variants in other proteins in the HDR pathway that are also hereditary breast and ovarian cancer tumor suppressors, such as BRCA2^28–30^ and BARD1^31,32^. As genetic testing for cancer risk becomes more common and additional genes are added to testing panels, the number of rare missense variants that inevitably become VUS will continue to increase. We anticipate that multiplexed functional assays can be used to functionally characterize such variants at scale, even before the variants are observed in results from genetic testing.

## Acknowledgements

We thank Jason Underwood and Katy Munson of the University of Washington PacBio Sequencing Services for assistance with long-read sequencing, Ronald Hause and Alan Rubin for helpful suggestions regarding statistical methods, Martin Kircher for help with analysis of long read sequences, Ethan Ahler for assistance with figures and the OSU Comprehensive Cancer Center Analytical Cytometry Shared Resource for sorting of the GFP-positive cells.

## Contributions

L.M.S. and J.G. created, barcoded and assembled the plasmid libraries for the expression of mutant BRCA1. L.M.S. amplified barcodes, sequenced and analyzed results of sorting experiments. M.M.I. established the HeLa-derived cell lines with integrated libraries and with T.B. and A.I.A., performed multiplexed HDR assay, sorted cells and extracted genomic DNA. S.F., J.S., and J.D.P. provided guidance. L.M.S., S.F., J.S., and J.D.P. wrote the manuscript.

## Methods

All enzymes unless specifically mentioned were purchased from New England Biolabs. Primer sequences can be found in Supplementary Table 3.

### Creating the HeLa DR-FRT cell line

A description of the HeLa-DR cell line can be found here^13^. To create a FLP-in version of HeLa DR, we stably integrated into the cells a flipase recognition target (FRT) sequence using the pFRT/lacZeo plasmid (Thermo Fisher). Zeocin resistant clones that had a single integration site detected by Southern blot were tested for high activity integration sites using the mammalian β-galactosidase activity assay (Gal-Screen, ThermoFisher). Clonal expansion of the selected colony established the HeLa DR-FRT cell line.

### Site-saturation mutagenesis libraries and barcoding

The HA-tagged BRCA1 N-terminal HindIII-EcoRI fragment containing amino acids 1-302 was cloned into the pUC18 plasmid. Three site-saturation mutagenesis libraries of BRCA1 were constructed using a previously reported inverse PCR-based method^19^. Pool1 has amino acid substitutions in amino acids 2-96, pool2 97-192 and pool3 193-302. For each codon, 30 base, mutagenic primers were ordered with machine-mixed NNK bases at the 5’ end of the sense oligonucleotide (N = ACTG, K = GT). The mutagenized HA-BRCA1 2-302 fragments were ligated into the EcoRI and HindIII sites of the BRCA1 cDNA in pcDNA5/FRT-TO vector that had been modified to remove a second EcoRI site in the gene for hygromycin resistance and a second multiple cloning site added at the MluI/NruI sites for the barcode. A 16 base, degenerate barcode encoded on oligos (pc5_barcode_longer_ W and _C) that had been annealed, extended, and digested with NotI/SbfI was then ligated into the second multiple cloning site. There were ~25,000 barcoded clones for each of the pool1 and pool2 libraries and ~50,000 for pool3 as assessed by colony forming units after barcode ligation and transformation. Metrics for the variant library cloning steps can be found in Supplementary Table 1.

### Assigning barcodes to variants using PacBio long reads

To prepare the circular SMRT-bell templates for the pool 1 and 2 variable regions and barcode, the intervening sequence between the barcode and the BRCA1 N-terminal variable region was removed by NotI/HindIII restriction digest, followed by end-repair and blunt-end ligation^33^. The ligations were transformed into E. coli to remove concatamers. The plasmids were then cut with SbfI and EcoRI to release the barcode and BRCA1 N-terminal variable region. For pool3 the entire SbfI and EcoRI fragment including the extra 2 Kb of sequences was released. Custom SMRT bell adapters pb_SbfI and pb_EcoRI were sticky-end ligated to the purified fragment. To make a working stock of 20 μM SMRT bell adaptors in 10 mM Tris, 0.1 mM EDTA, 100 mM NaCl, they were heated to 85°C and snap cooled on ice. The ligation reaction contained 0.3 pmol purified fragment, 0.4 μM of each adaptor, 0.25 μL of EcoRI-HF, 0.25 μL of SbfI-HF, 1X ligase buffer, and 0.5 μL of T4 ligase in a 25 μL reaction. The ligation was performed at room temperature for 30 minutes, then heat inactivated at 65°C for 20 minutes. To cut destroy SMRT-bells with the remaining plasmid backbone, 0.25 μL each XhoI and NdeI were added and incubated 15 min 37°C. And finally, to digest noncircular DNA, 0.5 μL each of ExoIII (Enzymatics) and ExoVII were added and incubated at 37°C for 15 minutes. The final SMRT bell fragments were purified via AmpurePB (Pacific Biosciences) at 1.8X concentration, washed in 70% ethanol, eluted in 15 μL 10mM Tris pH 8 and quantified by BioAnalyzer (Agilent).

Each BRCA1 library was sequenced on four SMRT cells on a Pacific Biosciences RS II sequencer. Barcodes and variable regions were identified as in ref. 34 as follows: Base call files were converted from the bax format to the bam format using bax2bam (version 0.0.2) and then bam files for each library from separate lanes were concatenated. Consensus sequences for each sequenced molecule in every library were determined using the Circular Consensus Sequencing algorithm (version 2.0.0) with default conditions (ccs and bax2bam can found in the PacBio Github repository, https://github.com/PacificBiosciences/unanimity/blob/master/doc/PBCCS.md). Each resulting consensus sequence was then aligned to a BRCA1 reference sequence using Burrows-Wheeler Aligner^35^ (http://bio-bwa.sourceforge.net/). Barcodes and insert sequences were extracted from each alignment using custom scripts that parsed the CIGAR and MD strings. For barcodes sequenced more than once, if barcode-variant sequences differed, the barcode was assigned to the variant that represented more than 50% of the sequences. Barcodes lacking a majority variant sequence were assigned the variant sequence with the highest average quality score as determined by the ccs algorithm. The barcode-variant extraction and barcode unification scripts can be found at https://github.com/shendurelab/AssemblyByPacBio/. Pool1 had 19,809 barcodes assigned to variants with 0 or 1 amino substitution encoding 1602 unique protein variants. Pool2 had 17,635 barcodes assigned to variants with 0 or 1 amino substitution encoding 1695 unique protein variants. Pool3 had 11857 barcodes assigned to variants with 0 or 1 amino substitution encoding 1987 unique protein variants. Additional metrics regarding the sequence processing for the barcode-variant assignments can be found in Supplementary Table 1. For all three BRCA1 libraries, a barcode-variant map file was created that contains each barcode and its nucleotide sequence.

### Integration of libraries into cells

For each of the three BRCA1 plasmid libraries (pools 1-3), 70-80 million HeLa DR-FRT cells in ten 10 cm tissue culture plates were transfected with 200 µg of pOG44 to express a modified flipase enzyme and 100 µg pcDNA5/FRT BRCA1 variant library. Plasmids are diluted in 10 mL Opti-MEM and incubated for 5 minutes. 300 µl of Lipofectamine 2000 (ThermoFisher) is diluted in 10 mL Opti-MEM for 5 minutes. The lipofectamine and plasmid dilutions are then combined and incubated for 20 minutes. The mixture was then applied directly to cells. After 24 hours, cells were trypsinized and transferred to ten 15 cm tissue culture dishes. Since the pOG44 flipase has reduced activity at 37°C, four to eight hours post transfer, the cells were moved from a 37°C humidified incubator to a 30°C humidified incubator for 24 hours. The cells were then returned to 37°C for an additional 24 hours. Approximately 72 hours after the initial transfection, the cells were trypsinized and transferred to twenty 15 cm plates containing selection media (50% fresh DMEM supplemented with 10% fetal bovine serum, 50% filter sterilized conditioned media and hygromycin B at 550 µg/mL). Hygromycin resistant cells are selected at 37°C for 24 hours. The cells are washed with sterile phosphate buffered saline (PBS) and the selection media is replaced after the first 24 hours and again every 48 hours until cell colonies are visible without a microscope (about 14 days). Colonies were then counted, trypsinized, resuspended in 20 mL culture media, and mixed thoroughly, colony counts can be found in Supplementary Table 1. 15 ml of the resuspended cells were frozen in 1 mL aliquots. 5 mL of resuspended colony mixture was plated onto three 15 cm plates (3 mL, 1.5 mL, 0.5 mL) and incubated for 24 hrs. The plate that was closest to a confluent monolayer of cells was passaged. The cells were passaged for an additional two weeks before performing HDR reporter experiments to assure loss of the unintegrated BRCA1 expression plasmid.

### HDR reporter assays, sorting, gDNA prep

HDR reporter assays and FACS sorts were performed for each of the three BRCA1 variant libraries in quadruplicate. A confluent 10 cm plate of HeLa BRCA1 variant cell line was trypsinized and resuspended in 10 mL of culture media. 65 μl of the suspension was plated in each of 48 wells across two 24-well tissue culture plates. 24 hours later, each well was transfected with 30 pmol of siRNA and 1.5 μl Oligofectamine (ThermoFisher). Oligofectamine was diluted with 6 μl Opti-MEM and siRNA was diluted with 25 μl Opti-MEM for 5 minutes. The dilutions were then combined and incubated for an additional 30 minutes. The transfection mixture was then applied directly to the cells. 24 hours later, the cells were trypsinized and transferred to four 6-well tissue culture plates, then incubated for another 24 hours. Each well was then transfected with 50 pmol of siRNA, 3 μg pCBASceI (for I-SceI expression), and 3 μl Lipofectamine 2000. Plasmid and siRNA were diluted in 125 μl Opti-MEM and Lipofectamine was diluted in 125 μl Opti-MEM, then incubated for 5 minutes. The dilutions were combined and incubated for an additional 20 minutes. The transfection mixture was then applied directly to cells. Four to six hours later, the culture media was replaced with fresh media. BRCA1 3’UTR siRNA, BRCA1 coding sequence siRNA and control siRNA were used in both rounds of transfection. After 24 hours, one well of cells treated with each condition was analyzed for GFP expression using flow cytometry to confirm transfection efficiency. If cells treated with control siRNA and 3’UTR siRNA fell within 7-9% and 4-7% GFP+ cells respectively, and cells treated with BRCA1 coding sequence siRNA were 1-2% GFP+, the experiment would proceed. 72 hours post-transfection, the cells were pooled according to treatment and sorted using fluorescent activated cell sorting (FACS) using an Aria IIu instrument. Cells were resuspended and pooled in filter-sterilized sorting buffer containing 1X Phosphate Buffered Saline (Ca^2+^/Mg^2+^ free), 5 mM EDTA, 25 mM HEPES pH 7.0, and 1% heat-inactivated fetal bovine serum dialyzed against Ca^2+^/Mg^2+^ PBS. A minimum of 500,000 GFP+ cells and a maximum of 2 million GFP-cells were collected per pool (Supplementary Table 2). Genomic DNA (gDNA) was extracted from the GFP+ and GFP-cells with a DNeasy Blood & Tissue Kit according to manufacturer instructions (Qiagen). DNAs were eluted in 200 μl Buffer EB.

### Barcode amplification and sequencing

To amplify the barcode from gDNA from the GFP-positive and negative populations were spread over 8 - 16 reactions containing 250 ug of gDNA each (220K – 440K genome equivalents by weight given HeLa triploid genome ~9 pg, see Supplementary Table 2). Reactions also contained Kapa2G Robust Polymerase (Kapa Biosystems), a primer that annealed to the SV40 promoter 5’ of the integrated plasmid and a primer adjacent to the barcode (SV40_F and newpc5bc_nexteraR). PCR was performed using the following conditions [95°, 5 min; {95°, 40s, 65°, 30s, 72°, 3 min} x ~28 cycles; 72°, 10 min]. The reactions produce a ~3,700 base amplicon specifically from integrated plasmids. The reactions for each sample were combined and the amplicons were purified by 0.5X Ampure and eluted with 10 mM Tris at 10% of the original reaction volume. 10% of the eluted PCR volume was re-amplified with primers containing sample indexes and Illumina cluster generating sequences (pc5bc_p5_F, nextIndex). Reactions also contained Kapa2G Robust Polymerase and 0.5X SYBR green II (ThermoFisher). PCR reactions are monitored on a Mini-opticon qPCR machine (Bio-Rad) and removed during exponential amplification using the following conditions [95°, 3 min; {95°, 20s, 65°, 30s, 72°, 20s} x 5-10 cycles]. The PCR reaction produced a 350-base amplicon that was purified using a double-Ampure protocol. First the large DNA fragments were removed by precipitation by addition of 0.6X volume Ampure beads to the reactions. Then the 350-base amplicons were purified from the supernatant using 0.9X volume Ampure beads. The samples were multiplexed and the barcodes and sample indexes were sequenced (single read, pcDNA5_barcodeSeq_F) on a Nextseq 500 High Output 75 base kit, reads per experiment can be found in Supplementary Table 2. For the pilot 16-plex experiment without barcodes, the region of BRCA1 containing the variants was amplified and directly sequenced.

### Variant scoring, classifications and depletion score

FASTQ files containing either barcodes for each sample and the barcode-map for each library were used as input for the software package Enrich2^22^. Enrich2 was used to count the barcodes, associate each barcode with a nucleotide variant, and then translate and count both the unique-nucleotide and unique-amino acid variants. Barcodes assigned to variants containing insertions or deletions were removed from analysis. The counts for each protein variant were converted to frequencies by dividing by the total number of variant counts for each sample. The ratio of the frequency of each variant in the GFP positive population over its frequency in the GFP negative population was calculated. That ratio for each variant was then normalized to the GFP+/GFP-ratio for the wild-type sequence at ln(GFP+/GFP-) = 0. Variants with multiple amino acid substitutions were removed from further analysis.

To construct the binary, functional / nonfunctional classifier for variants, we first determined that the standard deviation of the score decays according to read count in a way that can be modeled by a log_10_-log_10_ curve. We then modeled the decay of the standard deviation of scores from the control siRNA experiment. Next, we applied that model to the standard deviation of scores from the BRCA1 siRNA experiment and calculated a p-value and an FDR-adjusted q-value for each variant to determine if it was similar to the control siRNA experiment (q > 0.05) or significantly different from control experiment (q <0.05). Finally, we determined where the false positives in the control experiments occurred along the continuum of read counts to assign a read-count threshold for the BRCA1 siRNA condition. A read-count threshold is usually part of the heuristic applied to remove noise due to stochastic dropout from deep mutational scanning data ^16^. We removed variants with below that read count threshold from further analyses. These analyses are performed using R studio, an R markdown file containing all data manipulations is supplied as Supplementary File 1.

The depletion score represents the number of replicates in which a variant was found to be depleted from the BRCA1 siRNA GFP-positive population. Only variants that had been present above the read-count threshold in at least three replicates have depletion scores. Depletion scores for those 1699 variants can be found in Supplementary Table 5.

## Website accessions

ClinVar, accessed June 2017, minimum 1-star interpretation.

## Supplementary Materials

**Supplementary Figure 1.**
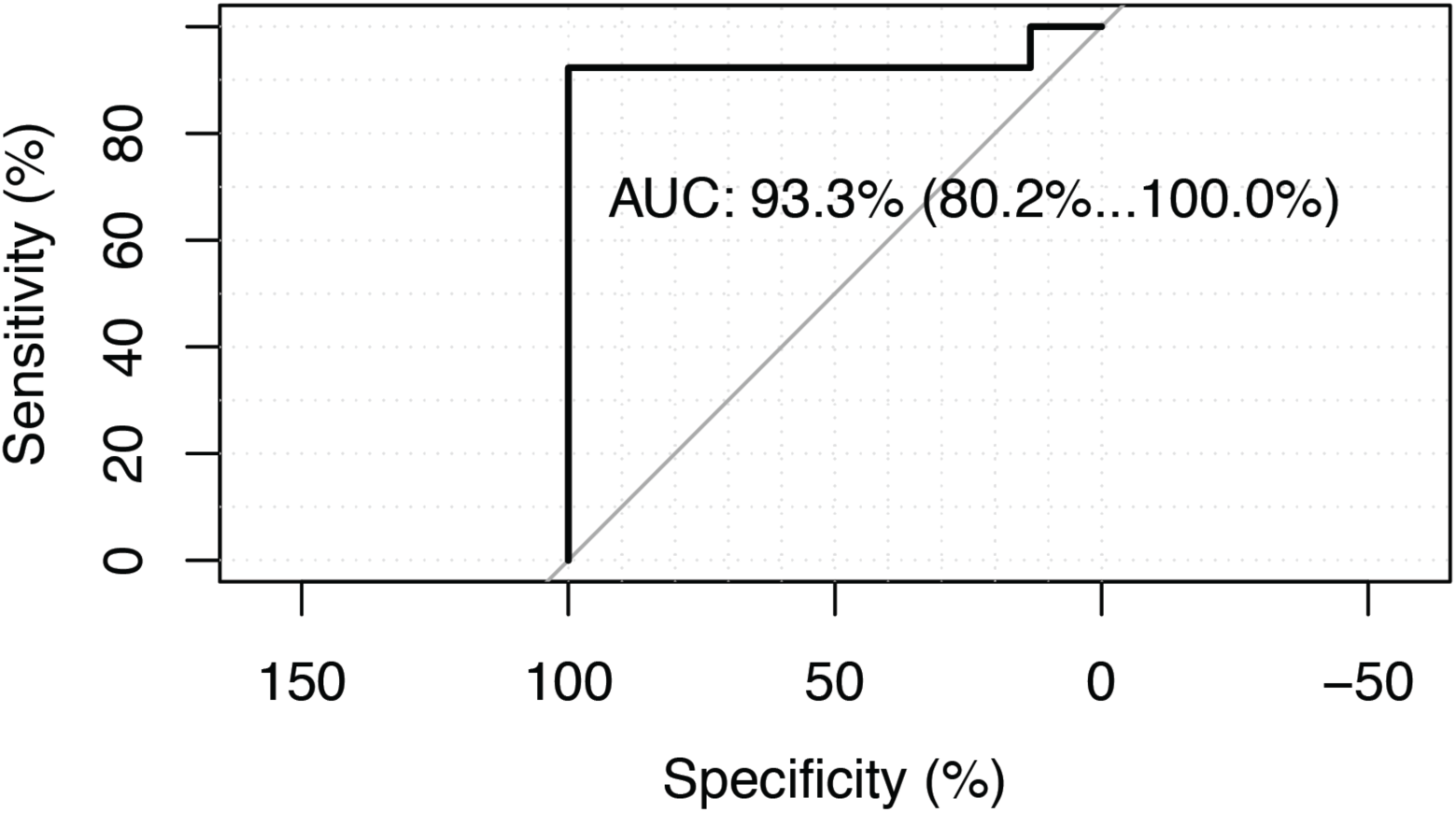
Receiver-Operator Characteristic curve for singleton HDR reporter assay results. N = 43, 30 known benign and 13 known pathogenic BRCA1 variants stratified by results from single HDR reporter assays from refs 1–4. Plotted with the pROC r package^5^. 90% confidence interval reported.

**Supplementary Figure 2.**
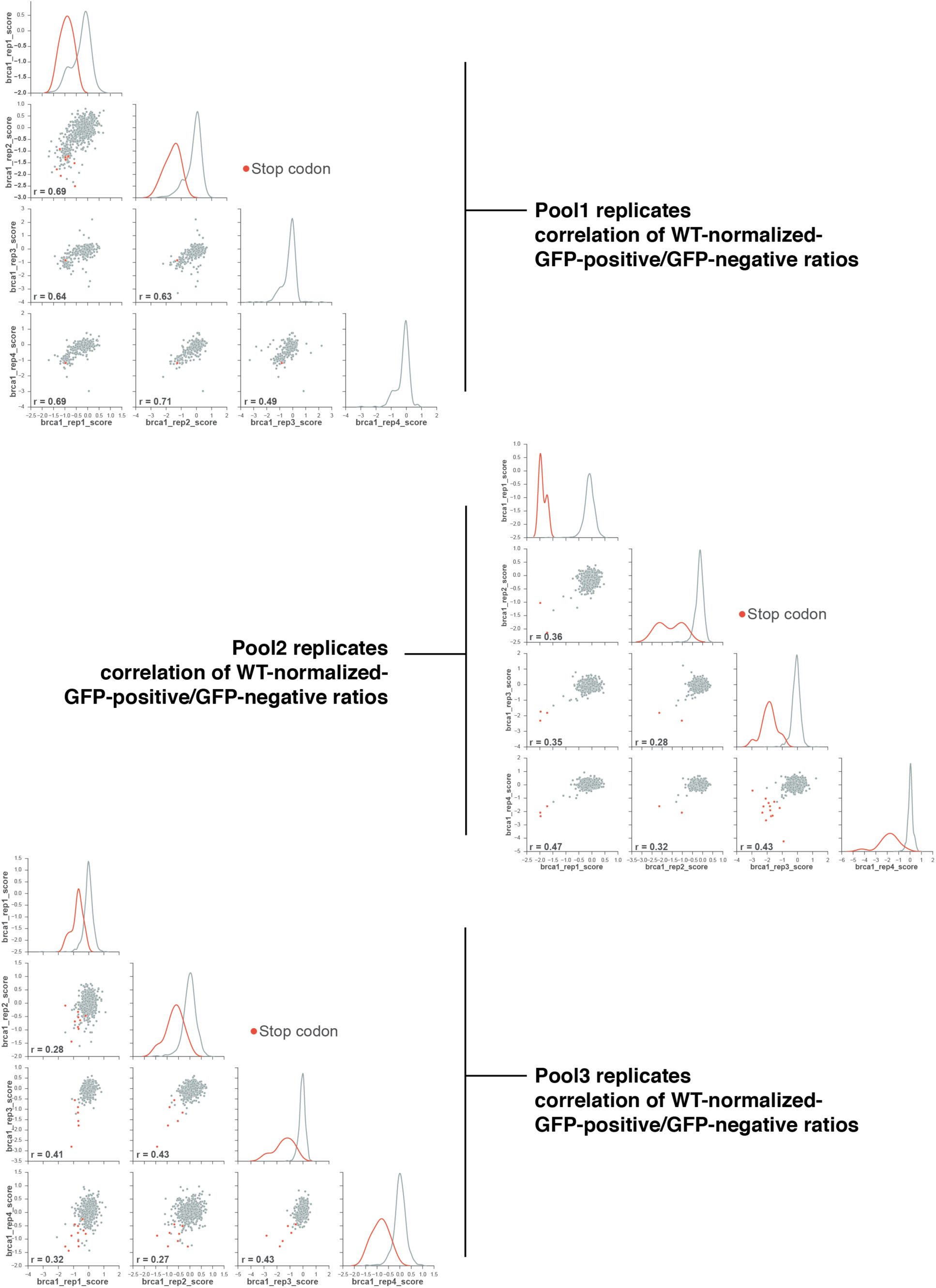
Scatter plots of multiplex HDR reporter assay scores from replicate experiments. Only variants above the read count threshold are included. Missense variants are grey and nonsense variants are orange. Pearson’s r reported.

**Supplementary Figure 3.**
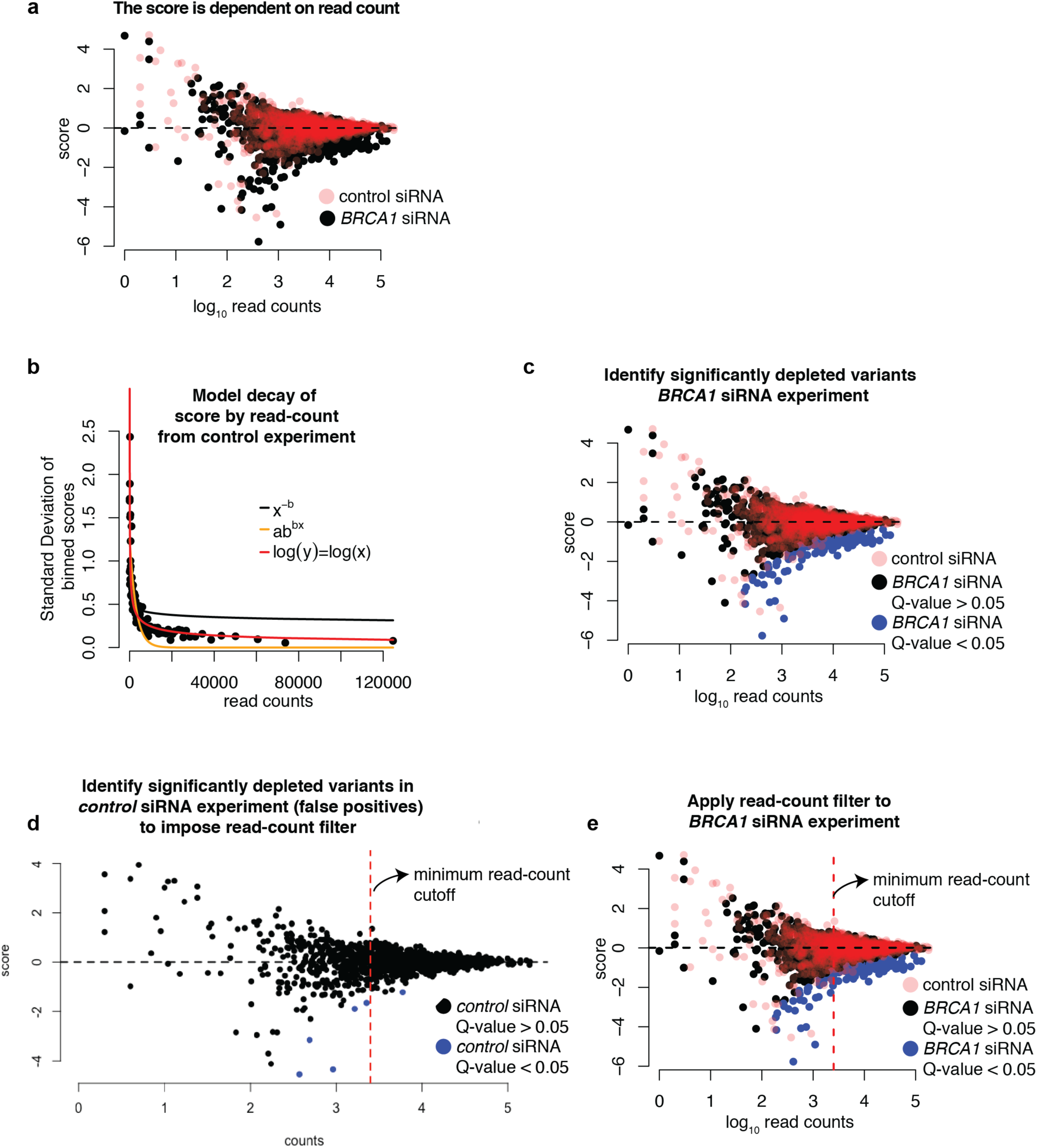
Illustration of the depletion classifier and read count thresholds using pool 1, replicate 1. **a,** The log of the WT-normalized, GFP-positive:GFP-negative ratios are on the y-axis and log_10_ read counts are on the x-axis for a single replicate of the HDR-reporter assay for codons 2-96. Variants from the control (red) or BRCA1 (black) siRNA conditions are indicated. **b,** Log(x) = log(y) best fits the decay of the standard deviation of the score in 100 variant bins, given the read counts in the control siRNA condition. Lines representing each model are indicated. **c,** The same plot as in (a) with variants significantly depleted from the GFP-positive population in the *BRCA1* siRNA condition, q < 0.05, colored blue. **d,** The variants in the control siRNA population (here, colored black) with the false positive (q < 0.05) colored blue. The dashed line represents the read-count threshold chosen to minimize the number of false positives. **e**, The same plot as in (a) with variants significantly depleted from the GFP-positive population in the *BRCA1* siRNA condition, q < 0.05, colored blue and the dashed line (from d) represents the read-count threshold. All R code and plots for the remaining pools and replicates can be found in the R Markdown supplied as supplementaryFile1.Rmd. Figures can be remade using the markdown file and data provided in SupplementaryTable4.tsv

**Supplementary Table 1.** Metrics for BRCA1 barcoded variant library construction, cell integration, barcode assignments and library quality. aa = amino acid, CFU = colony forming units

| library | pool1 | pool2 | pool3 |
| --- | --- | --- | --- |
| BRCA1 variable region aa positions | 2-96 | 97-192 | 192-302 |
| inverse PCR clones | 3.2 M | 1.6 M | 360 K |
| pcDNA5 cloning CFU | 180 K | 80 K | 140 K |
| barcode cloning CFU | 25 K | 25 K | 50 K |
| HeLa HDR FRT integrations | 54 K | 150 K | 35 K |
| PacBio SMRT cells | 4 | 4 | 4 |
| PacBio reads with more than 3 passes | 141720 | 162351 | 78875 |
| passed alignment filters (min mapQ 30, min quality score per base = 20, no softclipping, correct size barcode) | 61906 | 73254 |  |
| passed alignment filters (min mapQ 30, min quality score per base = 0, no softclipping, correct size barcode) |  |  | 54001 |
| unique barcodes | 27433 | 24497 | 26200 |
| barcode had one consensus read | 10963 | 6632 | 11945 |
| barcode had two consensus reads | 7484 | 5710 | 6259 |
| barcode had three consensus reads | 4283 | 4320 | 2669 |
| barcode had > 3 consensus reads | 4703 | 7835 | 5327 |
| all reads associated with a barcode were identical | 15966 | 17266 | 3193 |
| read assigned by major allele | 363 | 488 | 1836 |
| read assigned by highest quality score tie breaker | 141 | 111 | 5471 |
| barcodes associated with WT or single amino acid substitution | 19809 | 17635 | 11857 |
| barcodes associated with indel | 6689 | 6046 | 13426 |
| barcodes associated with >1 aa substitution | 935 | 816 | 917 |
| Unique variants with 0 or 1 aa substitution | 1602 | 1695 | 1987 |

**Supplementary Table 2.**
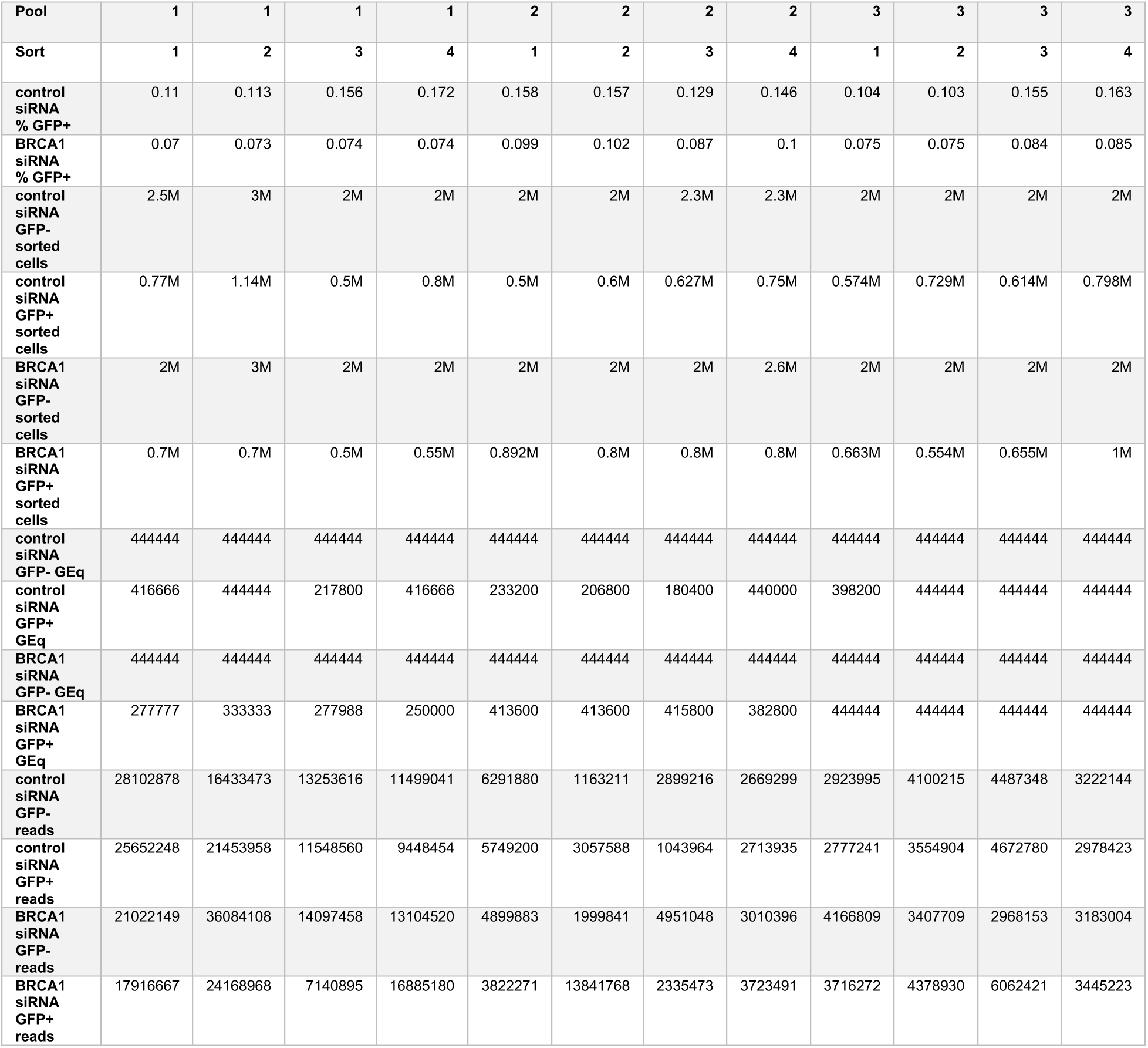
Metrics for FACS sorts, PCR amplification and DNA sequencing for replicate multiplexed HDR reporter assays. GEq = genome equivalents (9 pg per triploid HeLa genome)

**Supplementary Table 3.**
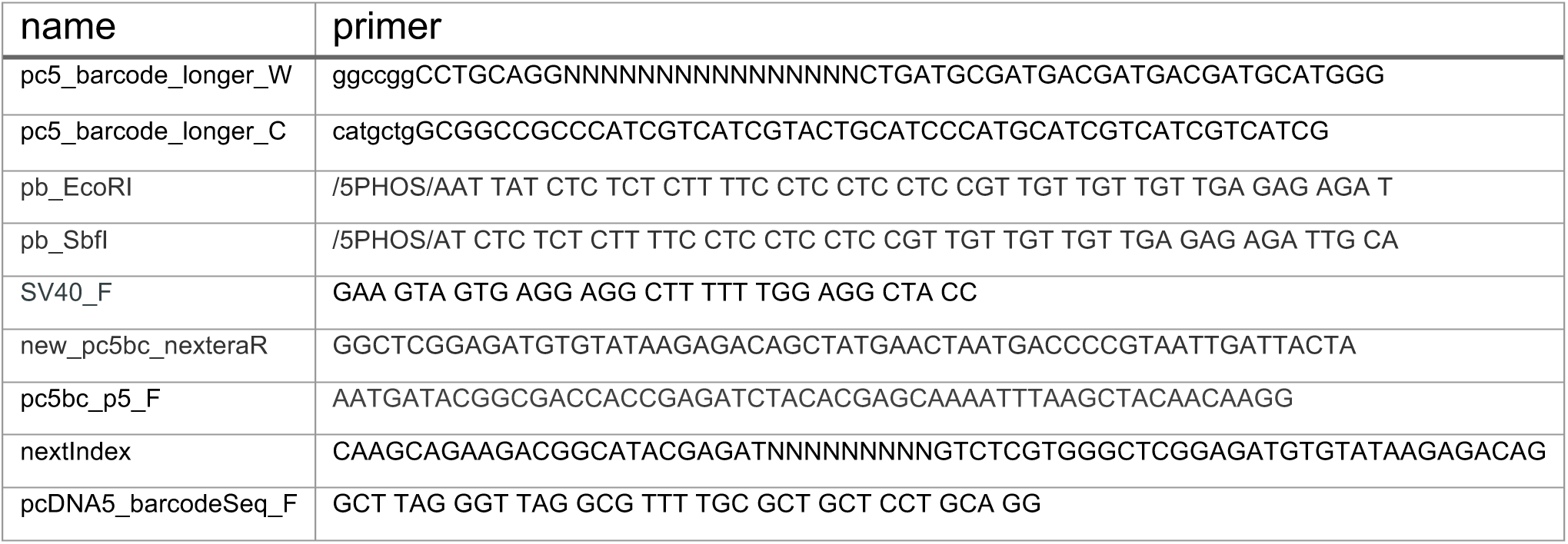
Primer sequences used for this study.

**Supplementary Table 4. Counts, scores and q - values for all variants in all replicates.**

**Counts:** Column headers have the name of the siRNA treatment (brca1 or control), replicate (sort 1-4), and population (noGFP or GFP). Columns appended with _counts are raw sequencing counts for each variant.

**Score:** Column headers have the name of the siRNA treatment (brca1 or control), replicate (sort 1-4) and are appended with _score. The score calculated by taking the ratio of GFP/noGFP of the frequency of each variant in each population, normalized to that of WT, scores are reported as the natural log.

Pool: The pool represents which amino acids are varied, pool_1 2-96, pool_2 97-192, pool_3 = 192-302.

**Variant**: Protein variants are listed using HGVS nomenclature.

**ProtPos**: Amino acid position

**mut**: mutant amino acid

**WT**: Wild-type amino acid

**pval**: Column headers have the name of the siRNA treatment (brca1 or control), replicate (sort 1-4). Columns appended with _pval are the raw p-value describing the significance of the score difference between a variant in the BRCA1 population and variants in the control siRNA population at a given read count.

**qval**: Column headers have the name of the siRNA treatment (brca1 or control), replicate (sort 1-4). Columns appended with _qval are false discovery rate adjusted p-values^6^.

**Sigcol:** Column headers have the name of the siRNA treatment (brca1 or control), replicate (sort 1-4). Columns appended with _sigcol determine the color of the points for the variants. q < 0.05 = blue, the remainder are black.

**Supplementary Table 5 | Depletion scores for 1,700 BRCA1 variants.**

Depletion scores for the 1,699 variants that passed the read count threshold in 3 or 4 replicates.

**Variant**: Protein variants are listed using HGVS nomenclature.

**ProtPos**: Amino acid position

**mut**: mutant amino acid

**WT**: Wild-type amino acid

**variantID**: Wild-type amino acid concatenated with protPos and mut

**mHDR_repPass**: Number of replicates in which the variant passed the read count threshold, variants with only 3 or 4 are included.

**mHDR_depletionScore**: Number of replicates in which the variant was significantly depleted from the BRCA1 siRNA GFP-postive population (q < 0.05).

**clinvar_clinsig**: The ClinVar variant interpretation as of June, 2017.

**Parvin_HDR_average**: singleton HDR scores from refs. 1–4.

**HDR_function_cat**: singleton HDR functional category. HDR > 0.5 = high, HDR < 0.5 = low.

**Starita_Y2H_score**: Multiplexed yeast-two-hybrid scores from ref. 3.

**Starita_E3_score**: Multiplexed ubiquitin ligase scores from ref. 3.

**Starita_HDR_predict**: HDR predictions from ref. 3.

**Bouwman_class**: functional predictions from ref. 7. **SGE_functionScore_aveVar**: Saturation Genome Editing (SGE) function score averaged for SNVs that make the same protein variant (Findlay et al. enclosed).

**SGE_lowRNAcount:** The number of times an SNV at a the codon position causes a >75% reduction in RNA as measured by SGE (Findlay et al. enclosed).

